# *In situ* quantification of individual mRNA transcripts in melanocytes discloses gene regulation of relevance to speciation

**DOI:** 10.1101/332569

**Authors:** Chi-Chih Wu, Axel Klaesson, Julia Buskas, Petter Ranefall, Reza Mirzazadeh, Ola Söderberg, Jochen B. W. Wolf

## Abstract

Functional validation of candidate genes for adaptation and speciation remains challenging. We here exemplify the utility of a method quantifying individual mRNA transcripts in revealing the molecular basis of divergence in feather pigment synthesis during early-stage speciation in crows. Using a padlock probe assay combined with rolling circle amplification we quantified cell-type specific gene expression in the native, histological context of growing feather follicles. Expression of Tyrosinase related protein 1 (TYRP1), Solute carrier family 45 member 2 (SLC45A2) and Hematopoietic prostaglandin D synthase (HPGDS) was melanocyte-limited and significantly reduced in follicles from hooded crow explaining the substantially lower melanin content in grey vs. black feathers. The central upstream transcription factor Microphthalmia-associated transcription factor (MITF) only showed differential expression specific to melanocytes - a feature not captured by bulk RNA-seq. Overall, this study provides insight into the molecular basis of an evolutionary young transition in pigment synthesis, and demonstrates the power of histologically explicit, statistically substantiated single-cell gene expression quantification for functional genetic inference in natural populations.

## INTRODUCTION

Feathers are a key evolutionary innovation of the epidermis arising in the dinosaur lineage basal to the avian clade (1, 2). Millions of years of evolution have modified the development of epidermal appendages from scales to the complex morphological structure of a feather (**Figure 1**). Its basic bauplan has been repeatedly expanded to a variety of feather types ranging from downy feathers for insulation to pennaceous wing feathers enabling flight (3–5). By integrating light-absorbing, chemical compounds modulating feather coloration the avian plumage further provides ample opportunity for inter- and intraspecific signaling (6). Mediating between the natural and social environment feather coloration constitutes an important component of natural and sexual selection with implications for adaptation and speciation (7, 8). Coloration differences arising within short evolutionary timescales within and among populations are thus believed to play an active role in the initial steps of species divergence (9, 10).

**Figure 1:**
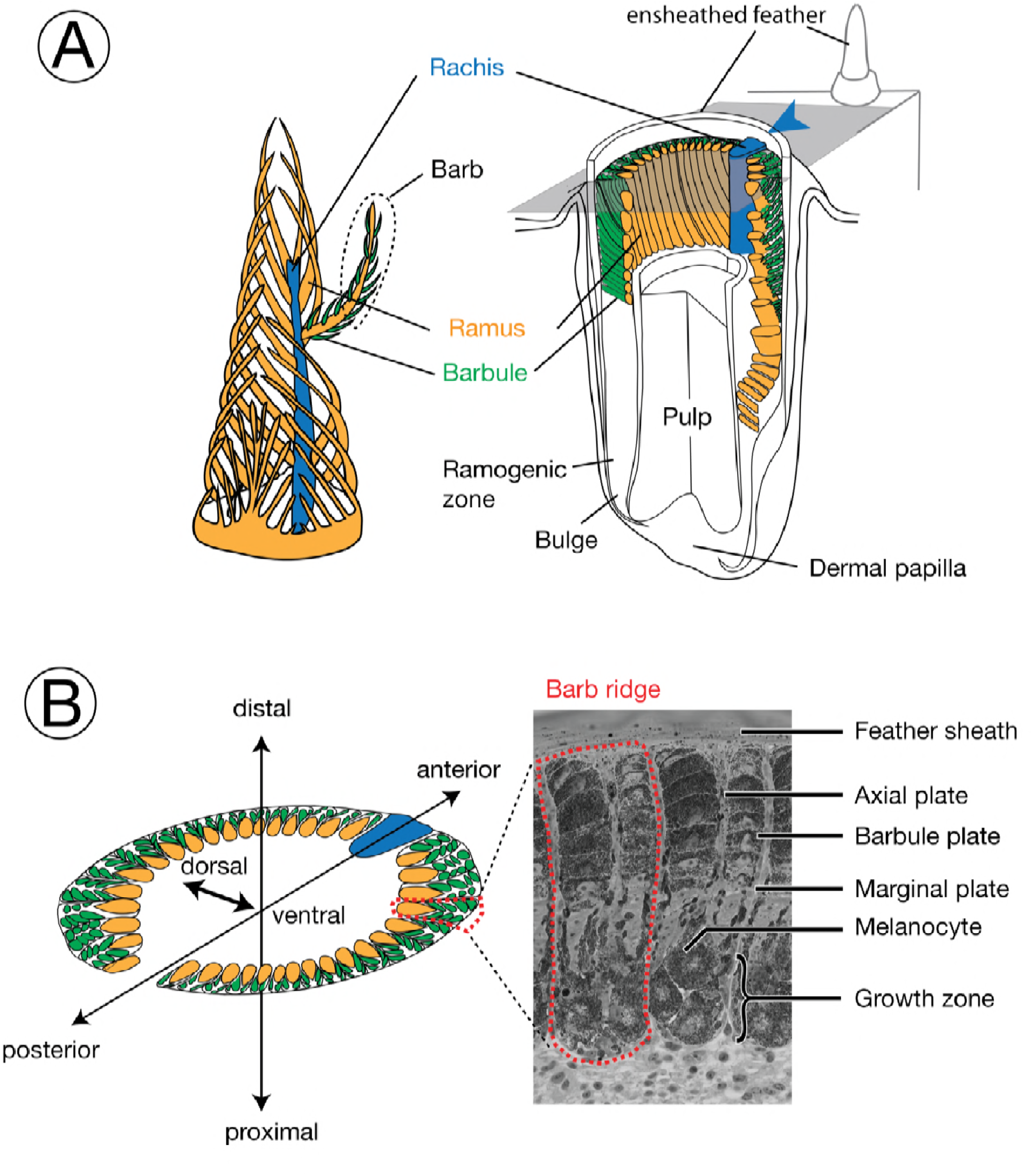
Schematic diagram and terminology of feather follicle morphology. The feather is a complex epidermal organ of cellular origin. Its basic bauplan follows a hierarchical structure of ramification into barbs and barbules. It develops within follicles, an epidermal modification derived from invagination into the dermis (53, 54). Cells fulfilling different functions in the mature barbs are derived from stem cells at the collar bulge of the basal layer (55). Developing progenitors of these cells migrate and gradually differentiate to form mature cells in barbs, including basal cells, marginal plate cells, axial plate cells, and barbulous plate cells (56). Development of the main feather progresses along the proximal-distal axis with helical displacement of barb loci. Barb loci emerge on the posterior midline and are gradually displaced towards the anterior midline where they fuse with the rachis. For more information on the feather morphology and development consult (17, 57). Colors define homologous structures in the respective panels; arrowheads indicating the rachis facilitate orientation in the remaining figures. **A) left:** Schematic, three-dimensional representation of a growing pennaceous feather with the afterfeather shown in front. **right:** Partial section a feather follicle. The cutting plane in grey shows the position of cross sections in panel B. **B) left:** Schematic cross section of a feather follicle with three-dimensional axes for orientation. **right:** Example of a cross-section from a growing, ensheathed carrion crow feather displaying two barb ridges.

The chemical compounds contributing to the vibrant colors of birds include metabolized ketocarotenoids producing yellow-orange color and porphyrins resulting in bright pink, yellow, red or green. Iridescence is mediated by refraction of light on feather microstructures often involving an ordered array of melanin granules within a keratin substrate (11). In this study, we focus on genesis and deposition of melanin polymers into the growing feather setting the basis for black, reddish or brown color space (6). While the molecular pathways of pigment synthesis and color patterning across the body are well characterized in mammals and insects (12, 13) inference in birds is usually indirect and often assumed to follow the general pathways characterized in other vertebrates (6) (but see 14, 15). Moreover, functional investigation is generally restricted to domesticated model species such as chicken (16, 17), duck (18), or quail (19, 20). Recently, however, high-throughput sequencing of genomes and transcriptomes has opened coloration genetic research to variation segregating in natural populations (21). Genome-wide association mapping approaches yield initial candidate genes contributing to variation in pigmentation within and among populations (22–24). Differential gene expression analysis between color morphs using bulk mRNA sequencing provides an independent axis to screen for genes potentially involved in color mediating pathways (25, 26). Yet, while useful to generate initial hypotheses, bulk mRNA sequencing integrates over a vast number of cells and cell types, and falls short of mapping gene expression relevant to the variance in pigmentation. Therefore, cell-type specific approaches measuring gene expression in the chromatophores themselves are required for functional characterization of candidate genes.

We here present a powerful approach allowing detailed quantification of transcript abundance in single cells. The basic approach is not limited to color genetics and can be used for any cell type and phenotype of interest. We focus on melanogenesis using an avian model of color-mediated early-stage speciation in European crows. Black carrion crows *Corvus (corone) corone* (CC) and grey-coated hooded crows *Corvus (corone) cornix* (HC) hybridize along narrow contact zones in Europe and Asia and show characteristic differences in the amount of melanin deposited in the plumage (**Figure 2A**). Evidence of assortative mating by phenotype and near-identical genomes differing by less then 100 differentially fixed base pairs (< 10^−6^ percent) make them a suitable model to investigate the genetic underpinnings of color divergence at an early stage of divergence (27). In previous studies, transcriptome analyses across several tissues revealed a limited number of differentially expressed genes between taxa. Differences were most pronounced in skin tissue with active feather follicles maturing into grey or black feathers. Among these differentially expressed genes, genes involved in melanogenesis were strongly enriched and predominantly down-regulated in the grey coated hooded crows. Together with population genomic scans for candidate genes that may causally be involved in divergence of the pigmentation pattern these results suggested divergence of one or a few upstream melanogenesis genes (26, 28). Yet, despite generalized differential expression throughout the melanogensis pathway, the transcription factor MITF with a central regulatory role in mammals showed no evidence of differential expression (26).

**Figure 2:**
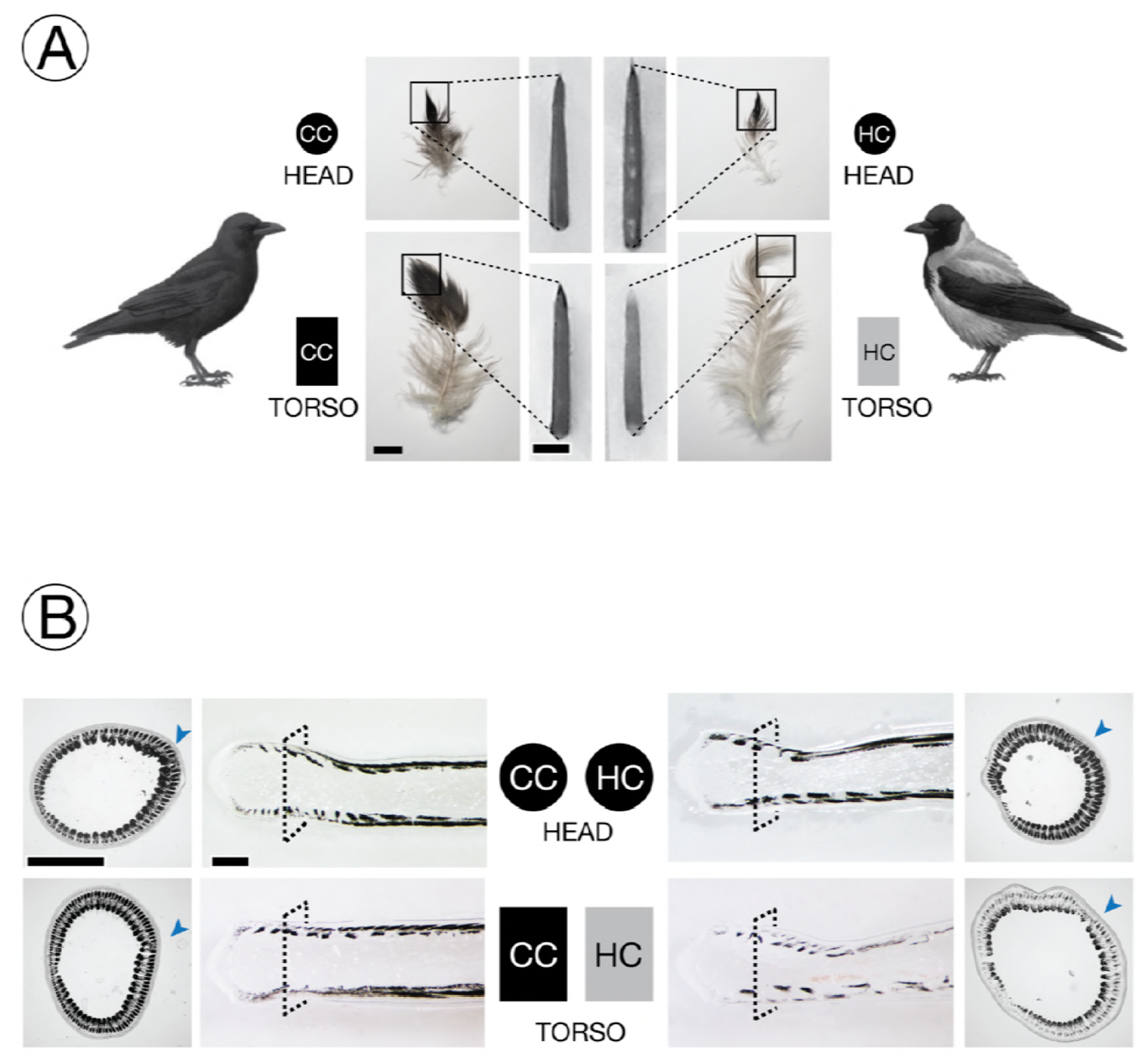
Experimental setup and phenotypic classification. A) Melanin-based plumage pigmentation differs between of all-black carrion crows (CC) and grey-coated hooded crows (HC). Growing feather follicles were sampled in the head region (circle) and on the ventral part of the torso (rectangle). Photographs adjacent to the schematic representation of birds show an example of semiplume, pennaceous body feather (bar = 1 cm) from torso or head for each taxon. Squares define the area of the mature feather represented by the ensheathed feather follicle (bar = 2 mm) which forms the raw material of the experiment. Lighter pigmentation of mature feathers of HC torso is already visible in ensheathed feather follicles (also see **Supplementary Figures S1 & S3**). Symbol color imitates the pigmentation of mature feather tips. [Bird drawings courtesy of Dan Zetterström]. **B)** Bright-**field** images from sections of ensheathed feather follicles. The dashed square on the longitudinal sections defines the corresponding position of the cross section at 1000 μm above the dermal papilla. The arrowhead in blue indicates the location of the rachis (see **Figure 1**). Dark areas represent regions where light-absorbing eumelanin is accumulated. Follicles from black-fathered regions show relatively higher eumelanin content in barb ridges than those sampled from the grey torso of hooded crows; bar = 0.5 mm.

Building upon information derived from bulk mRNAseq, we here characterize the molecular basis of (divergence in) avian melanin pigmentation in unprecedented detail. We first examined whether phenotypic and anatomical characteristics may contribute to explain the striking color contrast. We characterized melanocyte maturation, melanosome transport and the formation of barbed ridges in the native histological context of growing feather follicle of regenerating torso (grey in HC, black in CC) and head (black in both taxa) feathers from both taxa. We then quantified gene expression patterns of candidate genes both along the longitudinal axis of a feather and cross sections to reveal spatial-temporal expression patterns within and between taxa. To quantify mRNA transcripts in the cell type of origin we developed a histologically explicit padlock-probe assay coupled with rolling circle amplification allowing (RCA) (29) to partition variation in gene expression among taxa into its biologically relevant components.

Overall, this study provides detailed insight into the molecular mechanisms involved in feather melanization and gene regulatory evolution. It also demonstrates the power of the quantitative padlock assay allowing for statistical inference on gene expression differences in a cell-type specific context that can readily be transferred to other systems.

## MATERIAL AND METHODS

### Taxonomic considerations

The taxonomic status of the *Corvus (corone) ssp. s*pecies complex is contentious with some authors treating populations differing in plumage coloration as separate species (30). European all-black carrion crows and grey-coated hooded crows, however, have near identical genomes arguing against species status (28). Hence, until formal revision of taxonomic status recent work has treated carrion [*C*. (*corone*) *corone*] and hooded crows [*C*. (*corone*) c*ornix*] with uncertainty regarding taxonomic status (24, 28, 31). Given that for most part of the genome - including the target genes under investigation - genetic differentiation approaches null, taxa can essentially be treated as color morphs segregating within a population for the purpose of this manuscript.

### Population sampling

Crow hatchlings were obtained directly from the nest at an age of about three weeks. Hooded crows (*C.* (*corone*) *cornix*) were sampled in the area around Uppsala, Sweden (59°52’N, 17°38’E), carrion crows (*C.* (*corone*) *corone*) were obtained from the area of Konstanz, Germany (47°45N’, 9°10’E) (**Supplementary Table S1**). To avoid any confounding effects of relatedness, only a single individual was selected from each nest. After transfer of carrion crows to Sweden by airplane, all crows were hand-raised indoors at Tovetorp fieldstation, Sweden (58°56’55"N, 17° 8’49"E). When starting to feed by themselves they were released to large outdoor enclosures specifically constructed for the purpose. Carrion and hooded crows were raised and maintained under common garden conditions in groups of maximum six individuals separated by sub-species and sex.

Permission for sampling of wild carrion crows in Germany was granted by *Regierungspräsidium Freiburg* (Aktenzeichen 55-8852.15/05). Import into Sweden was registered with *Veterinäramt Konstanz* (Bescheinigungsnummer INTRA.DE.2014.0047502) and *Jordbruksverket* (Diarienummer 6.6.18-3037/14). Sampling permission in Sweden was granted by *Naturvårdsverket* (Dnr: NV-03432-14) and *Jordbruksverket* (Diarienummer 27-14). Animal husbandry and experimentation was authorized by *Jordbruksverket* (Diarienummer 5.2.18-3065/13, Diarienummer 27-14) and ethically approved by the European Research Council (ERCStG-336536).

### Feather sampling

For the purpose of the study, feather follicles needed to be sampled in a comparable, developmental stage with active expression of genes involved in the melanogenesis pathway. We therefore synchronized feather regrowth by plucking feathers on defined 3×3 cm patches on the torso (black in carrion crow, grey in hooded crow) and head (black in both subspecies) exploiting the natural disposition for molting during the months of July-September (32). Previous work has shown that regrown feathers are identical in color and shape to the original feathers (28). We restricted sample collection to the molting seasons in August of 2015 and 2016, corresponding to an age of 1 and 2 years, respectively (**Supplementary Table S1**). This excludes any potential ontogenetic changes in coloration sometimes occurring at the juvenile molt 4-6 weeks after fledging (32). We then allowed feathers to regrow for 10 to13 days, at which stage the previously plucked areas of skin contained densely spaced feather shafts with the first parts of feathers about to protrude from the shafts. Regenerating feather follicles ranging from 0.8 to 1.0 centimeters were collected in a stage of early development prior to apoptosis and keratinization (**Supplementary Figure S2 & S3**). Immediately after retrieval follicles were fixed in phosphate buffered saline (PBS) solution containing 4 % paraformaldehyde (AH Diagnostics, Sweden). This material forms the basis of the subsequent histological examination and mRNA quantification (29).

### Histology of melanocytes and characteristics of melanosomes

Feather follicles pretreated with paraformaldehyde (PFA) as described above were dehydrated in an ethanol serial dilution of 70%, 95%, and 99.9%, and then embedded in Technovit 7100 (EMS, USA). Serial plastic cross-sections of 3 μm in thickness were obtained using a microtome (MICROM HM 360, ThermoFisher Scientific Inc., USA), and subsequently stained with hematoxylin (Merck & Co. Inc., USA) and eosin (BDH Chemicals, UK). To identify the morphologic features of melanosomes, PFA-fixed feathers were dissected and squeezed, and then wet-mounted with PBS. Bright-field histological or whole-mounted images were taken using a Leica Imager M2 equipped with a Hamamatsu digital camera C1440.

### Quantification of targeted mRNA molecules in a single-cell context

#### In situ cDNA synthesis, hybridization of padlock probes and rolling circle amplification

The abundance of mRNA transcripts was quantified *in situ* for four gene targets, HPGDS, SLC45A2, TYRP1, and MITF acting in different parts of the melanogensis pathway. For *in situ* quantification we followed Weibrecht et al. (2013) using padlock probes in combination with rolling circle amplification (RCA) (see **Figure S11**) on cryosections of PFA-fixed feather follicles which were infiltrated with a serial gradient of 15% and 30% sucrose in PBS, and then embedded in OTC Cryomount medium (Histolab AB, Sweden). The ACTB which shows no expression differences between taxa across several tissues including feather follicles (26) was included as the internal reference to control for technical variation in staining efficiency among sections.

First, appropriate sites for Locked Nucleic Acid (LNA^TM^) primers initiating cDNA synthesis and for padlock probes hybridizing to the cDNA were designed on gene model of the *Corvus (corone) cornix* reference assembly ASM738373v2 available at NCBI (https://www.ncbi.nlm.nih.gov/) for five targeted genes, including TYRP1 (XM_010392933), HPGDS (4 isoforms: XM_010411872, XM_010411873, XM_010411874, XM_010411875), SLC45A2 (2 isoforms: XM_010395148, XM_019283413), MITF (14 isoforms: XM_010390220, XM_010390221, XM_010390223, XM_010390224, XM_010390225, XM_19280403, XM_19280404, XM_19280405, XM_19280406, XM_19280407, XM_19280409, XM_19280410,

XM_19280411, XM_19280412), and ACTB (XM_010392369). Where several transcript isoforms were annotated constitutively expressed exons were targeted (see **Table SB** for primer and padlock probe design). To guarantee the same specificity across individuals, and importantly across taxa, primers were targeted at monomorphic regions devoid of genetic variation segregating within and between taxa (28). Using the Padlock Design Assistant version 1.6 (29) we first identified a unique, 150-bp DNA fragment comprising an probably even number of the four nucleotides that was suitable for both LNA primers and padlock probes. Blast searches at a statistical significance level of e^−4^ and e^−9^ of the resulting candidate sequences of LNA primers and padlock probes against the *Corvus (corone) cornix* genome (reference ASM738373v2), respectively, confirmed unique matching. One μM final LNA primers with approximately 20 bp in length and a GC content of approximately 50 % (**Supplementary Table S2**) initiated *in situ* reverse transcription by RevertAid H Minus Reverse Transcriptase (ThermoFisher Scientific Inc., USA) at the final concentration of 20 U/μl with 1U/ μl of Ribolock ribonuclease inhibitor (ThermoFisher Scientific Inc., USA) and 0.5 mM dNTP (29). The remaining mRNA was digested with RNase H (0.4U/μl) (ThermoFisher Scientific Inc., USA), and padlock probes were hybridized at both ends to the single-stranded, *in situ* cDNAs. Ligation of the gap remaining at the hybridized 3’ and 5’ ends of padlock probes with Tth Ligase (0.5U/μl) (Genecraft Inc., Germany) circularized the padlock probe preparing them for subsequent rolling circle amplification (RCA) that was used as a basis for fluorescent labeling of the *in situ* cDNA molecules. For the best *in situ* spatial signaling of RCA by Phi 29 polymerase (1U/μl) (ThermoFisher Scientific Inc., USA), the 3’-end of a LNA primer were designed to overlap with the hybridization site of the padlock probe by at least 6 bp (29). After RCA specific fluorescently labeled detection oligonucleotides were hybridized to detection sites of long strand of RCA products derived from each specific padlock probe. This results in highly localized fluorescence dot-signals representing original mRNA molecules (**Figure 3**). Further details are provided in the Supplementary Material.

**Figure 3:**
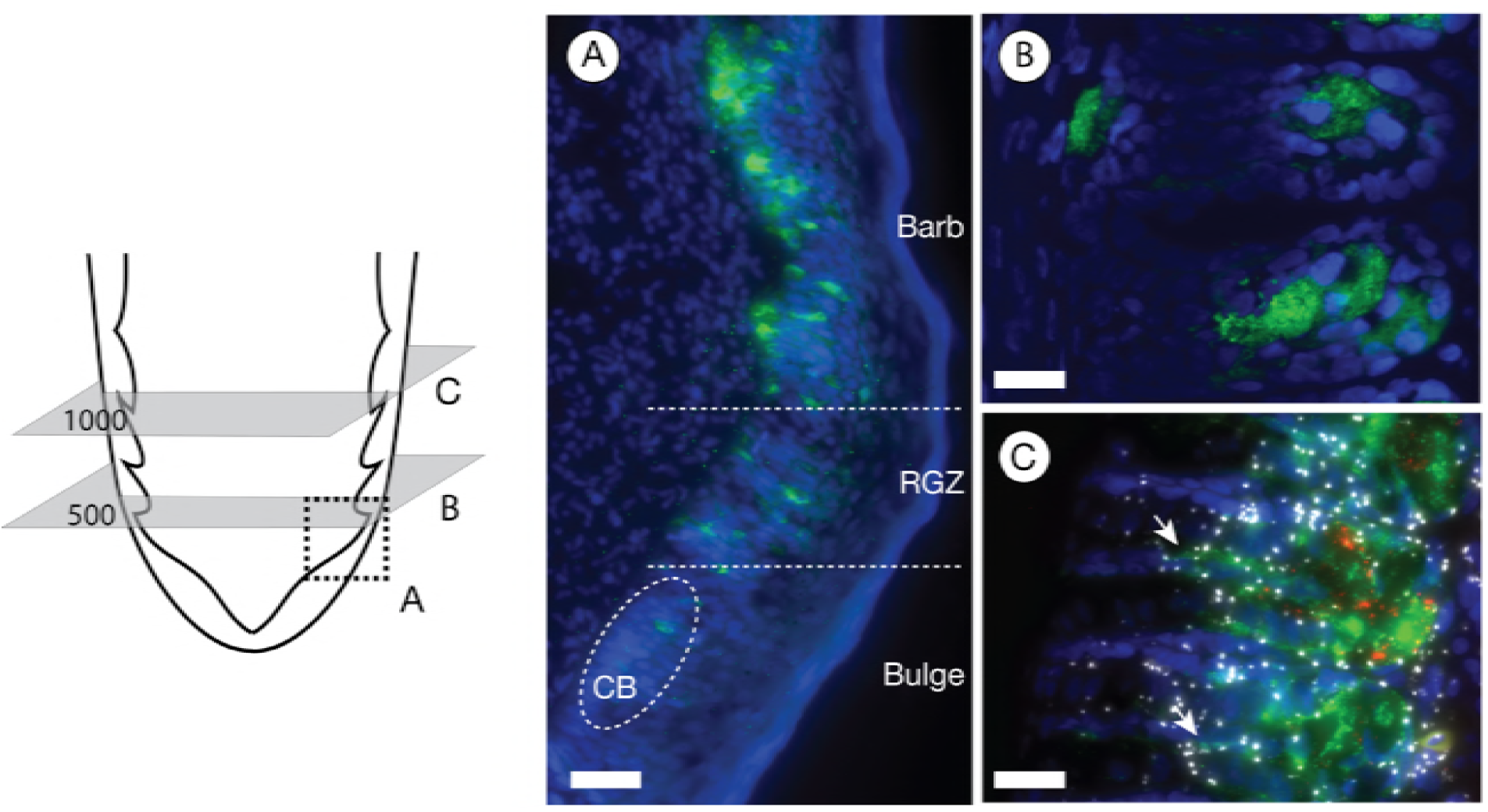
Melanocyte characterization. The schematic of the proximal end of a feather follicle on the left side depicts the cutting planes for the images to the right. Melanocytes are visualized by immunostaining against the TYRP1 protein (in green); nuclei were stained with Hoechst 33342 (in blue). **A)** The longitudinal section represented by the area of the dashed square illustrates the emergence of melanocytes in the collared bulge (CB) and their subsequent maturation in the ramogenic zone (RGZ) with increasing accumulation of the TYRP1 protein; bar = 50 μm. **B)** Cross section showing barb ridges at an early developmental stage at 500 μm above the dermal papilla. The TYRP1 protein begins accumulating in a tubular cell body located in the ventral growth zone; bar = 20 μm. **C)** Protein immunostaining combined with *in situ* mRNA padlock probe detection on a cross section at 1000 μm above the dermal papilla. At this developmental stage, melanocytes appear fully maturated: the TYRP1 protein (in green) is predominantly accumulated in a spherical cell body sending dendritic cytoplasm into barbules plates (white arrow). Signals of rolling circle amplification (RCA) of *in situ* mRNA padlock probing tagging individual TYRP1 mRNA transcripts (shown in red) are confined to the cell body. In contrast to melanocyte specific gene expression of TYRP1, mRNA of the internal control gene ACTB (in white) is ubiquitously expressed across other cell types; bar = 20 μm.

### Quantification of mRNA abundance along the proximal-distal axis of a feather

*In situ* mRNA abundance was quantified by counting independent, fluorescent signals on a Leica fluorescence microscope Imager M2 equipped with a Hamamatsu digital camera C11440 and corresponding filter setting. Fluorescent dyes were chosen such that mRNA abundance of three different genes could be investigated simultaneously on the same histological section. The first combination of detection oligonucleotides included the FITC fluorophore (filter excitation: 450 - 490 nm, filter emission: 500 - 550 nm) for the ACTB gene, the Cy3 fluorophore (filter excitation: 538 - 562 nm, filter emission: 570 - 640 nm) for SLC45A2 and the Cy5 fluorophore (filter excitation: 625 - 655 nm, filter emission: 665 - 715 nm) for HPGDS. The second set of detection oligonucleotides consisted of ACTB labeled with the FITC, MITF labeled with the Cy3, and TYRP1 labeled with the Cy5 (**Supplementary Table S2**). To investigate gene expression for both sets on the same histological section, padlock probes of the first gene set were stripped off with 20 % formamid at 50 °C for 30 minutes. Absence of any remaining fluorescence residues was confirmed prior to hybridization with detection oligonucleotides of the second gene set.

To capture all RCA *in situ* signals across the 10μm cryosection each section was scanned at different focal planes resulting in z-stacks of images. Prior to quantification each z-stack of images was algorithmically merged into a best-focal-point image using ImageJ ver. 1.51h (33) with the Stack Focuser plugin (34), optimally representing all RCA signals present in the sample. To ensure comparability between images taken separately for the control, dye set 1 and dye set 2, all images for the same section were aligned using a feature based alignment algorithm (35) in Matlab ver. 2016b. For some special cases, the alignment needed to be performed manually. This was done by using the ImageJ plugin Align Image by Line ROI (33). Quantification of signal position, intensity and cumulative number of RCA signals was automated using a Cellprofiler ver. 2.4.0rc1 pipeline (36). The basic steps of the pipeline were segmentation of the feather outline, detection of fluorescent signals, and detection of autofluorescence. The outline of the feather was extracted by enhancing the edges followed by morphological operations. The fluorescent signals were segmented using an adaptive local thresholding method based on ellipse fit (37). Regions with high autofluorescence were detected on images acquired before staining, and used to mask out the detected signals. All scripts are available in the supplement scripts and codes.

### Defining melanocyte boundaries

The abovementioned approach measures abundance of mRNA molecules for the entire image including cell types other than melanocytes. To specifically quantify mRNA abundance in melanocytes, we determined cell boundaries in cross sections using protein immunostaining against the TYPR1 protein. The TYRP1 protein is suited as a melanocyte-specific marker for the following reasons: (i) in humans the protein is exclusively found in skin (38); (ii) based on UniProtKB gene ontology data it is functionally limited to melanogenesis where it serves as a central downstream regulator of melanogenesis (humans: http://www.uniprot.org/uniprot/P17643, mouse: …/P07147, chicken: …/O57405) (39); (iii) in crows, TYRP1 transcripts are abundant only in skin with feather follicles (mean read count forebrain/liver/gonads/black feathered skin: 11/12/40/21,121) and (iv) within feather follicles the protein product is only detectable in melanocytes (28).

Immunostaining was performed with a primary antibody against human TRP1 at 1:500 dilution (ab83774, Abcam Co., USA) and secondary goat-anti-rabbit IgG H&L antibody (ab150077, Alexa Fluor 488, Abcam Co., USA) at 1:200 dilution. In addition, TYRP1 and ACTB mRNAs were visualized using the padlock-RCA approach described above, and cell nuclei were stained using 1 μg/μl Hoechst 33342 (ThermoFisher Scientific Inc., USA; filter excitation: 335 - 383 nm, filter emission: 420 - 470 nm). Combining the information from cell identity, melanocyte specific mRNA and protein expression (TYRP1) and ubiquitous mRNA signals (ACTB) melanocyte boundaries could be localized (**Figure 3, Supplementary Figure S7C**). Cell boundaries were determined on cross sections at 1000 μm distal to the dermal papilla for a subset of three follicle samples of carrion crows (**Supplementary Figure S7C**). Since the inferred cell body boundaries consistently coincided with the melanin cluster at the ventral end of the barbed ridges, melanin patterns could be used to extrapolate melanocyte boundaries in all other sections (**Supplementary Figure S7D**).

### Quantifying melanocyte-specificity of gene expression

To assess melanocyte-specificity of gene expression we quantified total mRNA abundance in five barb ridges for a total of 115 cross sections at both 500 μm and 1000

μm of each individual (torso HC, CC and head CC, HC) (**Supplementary Figure S7A**). In addition, we quantified the number of *in situ* tagged mRNA molecules residing exclusively in an area reflecting melanocyte boundaries delimited by TYRP1 protein immunostaining (**Supplementary Figure S7C**). Subtracting the latter from the total number of counts in barb ridges yields the number of transcripts expressed in cell types other than melanocytes (**Supplementary Figure S7D**). Normalizing both melanocyte-specific and unspecific counts by the respective area in the cross section provides a relative measure of transcript density, δ_mel_ and δ_other_ respectively. Since the expectation of a ratio of two random variables is generally not equal to the ratio of the expectation of the two random variables (40) we summed across all counts and normalized by the cumulative volume to obtain a single measure across sections for follicles from each of the eight combinations (CC/HC, torso/head, 500/1000 μm) Assuming similar levels of signal detectability in both regions allows defining a standardized measure of melanocyte-specificity as the S_mel_= δ_mel_ / (δ_mel_ + δ_other_). A value of 0.5 indicates equal density in both areas and hence ubiquitous gene expression independent of cell type. A value of 0.5> S_mel_ ≥1 indicates an excess of expression in melanocytes reaching 1 if expression were fully limited to melanocytes. Values of 0≤ S_mel_>0.5 indicate a depletion of gene expression in melanocytes relative to other cell types.

### Statistical comparison of transcript abundance

A major goal of this study was to quantify variation of *in situ* gene expression in melanocytes with respect to taxon (carrion/hooded crow), body parts (head/torso) and color (black/grey). To identify homologous regions for cross sections matched by developmental stage we first quantified transcript abundance along the feather using the padlock-RCA approach on longitudinal sections of the follicle (**Figure 6**). Two suitable regions were identified: a region of early differentiation at 500 μm distal of the dermal papilla, and a mature, differentiated region at 1000 μm where (i) expression levels of the central transcription factor MITF peaks and (ii) the lowly expressed SLC45A2 and HPGDS genes start saturating. (iii) Moreover, we found that 1000 μm constitutes a compromise between maximal gene expression levels and density of the melanin attenuating the signal of RCA products. At each region five serial cross cryosections of ten μm in thickness were sampled covering together approximately at least one whole melanocyte. For each section, accumulated RCA signals within melanocytes of five barbs located next to the rachis were measured and summed across the serial sections for each barb ridge as the process described above (**Supplementary Figure S7B, & D**). The sum of RCA signals of one barb ridge reflects the number of detected mRNA transcripts of an average of 3 (range: 2-4) melanocytes present per barb.

To control for technical differences in staining efficiency among and within sections, the abundance of each target gene was normalized relative to the ubiquitously expressed ACTB gene (an independent ACTB reference was used for each dye set, see above). Statistically, this can be modeled using the binomial distribution where RCA signal counts of the target gene *K* are expressed as a function of the total number of counts *N* of both target gene and ACTB. The probability of obtaining *K* successes is given by 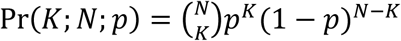. The probability of success *p* can here be thought of as the expression of the target gene relative to the ACTB control. A generalized linear mixed model with binomial error structure was then used to assess significant differences in relative gene expression (parameter *p*) across taxa (carrion/hooded crow), body area (head/torso) and color (black/grey). Due to over-dispersion in the models variation not accounted for by any of these explanatory factors arising by individual differences among sections was modeled as a random factor. A set of candidate models including all possible, additive combinations of explanatory factors was compared using the Akaike Information Criterion (AIC). To control for possible effects of melanocyte maturity the position of the cross-section was included in all models as a factor (500 μm vs. 1000 μm). Following Burnham and Anderson (41) models differing by ΔAIC ≤ 2 **were** considering to be supported by the data. Akaike’s weight wAIC was used to assess the relative importance of each explanatory variable. Statistical models were run for each target gene separately using the R package *lme4* of R version 3.4.1. (42, 43).

## RESULTS

### Morphological and anatomical characterization

Pennaceous wing and tail feathers, as well as body feathers covering the head and chest appear black in both carrion and hooded crows. Body feathers of hooded crows are light grey in contrast to all-black carrion crows (**Figure 2A, Supplementary Figure S1**). In both taxa semiplume body feathers are not uniformly pigmented. The intensity of eumelanin pigmentation increases from the proximal end, which is dominated by plumulacous barbs, to the fully pigmented, pennacoues tip of the feather. Feathers from hooded crow torso, however, are overall significantly lighter, but show the same pigmentation gradient along the feather (**Figure 2A, Supplementary Figure S1**). As the basis for all subsequent analyses we collected feather follicles from head (black in both taxa) and the ventral side of the torso (black in CC, grey in HC) at an early stage of development when still ensheathed by a protective outer epidermal layer (**Figure 1A & 2A, Supplementary Figure S2 & S3**). As feathers grow from proximal to distal, our sample represents the visible tip of the feather reflecting the natural contrast in plumage coloration between taxa in the torso region (**Figure 2A**).

Both longitudinal sections and cross-sections illustrate that differences in coloration are a direct consequence of taxon-specific variation in the level of melanization (**Figure 2B**). In both black and grey feathers melanin was deposited in rod-shaped melanosomes aggregating into higher-order granular structures (**Supplementary Figure S4**). While sections of black feathers (head CC & HC, torso CC) were saturated with melanin granules locally absorbing all light, grey feathers of hooded crow torso were more sparsely pigmented (**Figure 2B, Supplementary Figures S5 & S6**). Independent of the overall intensity, melanin was not uniformly distributed within barb ridges. Cross-sections illustrate that melanosomes were restricted to barbule plate cells separated by a melanin-free axial plate. Melanin was further concentrated at the dorsal end of barb ridges flanking the feather sheath, and ventrally in the growth zone adjacent to the ramus. The ‘melanin desert’ in between corresponds to rows of approximately three to four keratinocytes. In grey feathers, melanin levels appeared to be more depleted dorsally relative to the ventral growth zone of the barb ridge coinciding with the location of mature melanocytes (**Figures 1B & 2B, Supplementary Figure S5 & S6**).

Melanin is synthesized in melanocytes that are derived from activated progenitor cells at the very base of the follicle. Melanocyte maturation followed feather development in proximal-distal direction in both taxa: developing melanocytes first appeared in the collar bulge and were sparsely distributed in the basal layer of the ramogenic zone until about 500 μm from the distal end of the follicle (**Figure 3A**). At this stage most melanocytes were rod- to spindle-shaped (**Figure 3B**). At 1000 μm where barb ridges are fully formed, melanocytes were fully differentiated and spherical in shape (**Supplementary Movie S1**) measuring on average 32.8 ± 6.4 μm in diameter (n = 16). Mature melanocytes were situated in the ventral growth zone of a barb ridge and developed dendrite-like outgrowth of cytoplasm passing through cellular space in the axial plate and depositing melanosomes into keratinocytes of the barbule plates (**Figures 1B & 3C, Supplementary Movie S2**). Dendrite outgrowth appeared to be bounded by the marginal plate and axial plate, and thus limited within single barb ridges. Melanocyte development was accompanied by melanin deposition appearing in the middle-upper region bulge, the collar bulge, with a significant increase in the ramogenic zone (**Figure 3A**, **Supplementary Figure S7A**). In addition to the proximal-distal axis of feather maturation, the development of barbs also followed the anterior-posterior maturation pattern. Anterior barb ridges close to the rachis were developmentally more advanced than posterior barbs (**Figure 2B, Supplementary Figure S5, S6 & S7B**) as expected by the relative age of rami **(Figure 1A)**. Qualitatively, taxa did not differ in any of the morphological features considered above.

### Cell type specificity of gene expression

Next, we quantified gene expression for a set of four candidate genes that based on previous work 1) were involved in melanogensis, 2) covered a large range of expression levels and 3) were suggested to play a role in pigmentation divergence (see 26 and references therein). Gene targets included down-stream ‘effector’ genes SLC45A2 and TYRP1. SLC45A2 is a solute transporter of the basic melanin unit L-Tyrosine from the cytosol of melanocytes into the melanosome. The melanosomal TYRP1 enzyme catalyzes tyrosine metabolism and is thus directly involved in melanin synthesis. TYRP1 is predicted to be under control of MITF, a central regulatory transcription factor of the melanogensis pathway (44). HPGDS encodes the PGDS protein that may indirectly interact with MITF, but is generally not well characterized in the context of melanogensis (see discussion). Based on bulk mRNA sequencing of skin tissue of torso including regrowing feathers SLC45A2, TYRP1 and HPGDS showed significantly higher levels of steady state mRNA in carrion crows (26). The regulatory core gene MITF, however, was not differentially expressed. This may be due to a different role in avian regulatory networks or due to ubiquitous expression in cell types other than melanocytes diluting the effect.

To assess the functional role of these gene targets in avian melanogenesis we first quantified melanocyte-specificity of gene expression. TYRP1 mRNA transcripts almost exclusively accumulated in the spherical melanocyte cell body with only rare traces in dendrites, and were lacking altogether in other cell types (**Figure 3C, 4, Supplementary Figure S8**). Transcripts of HPGDS and SLC45A2 were also limited to the barb ridge growth zone likewise supporting melanocyte specificity (**Figure 4, Supplementary Figure S8**). Expression of MITF and the ACTB control gene, however, was not restricted to melanocytes. Transcripts of both genes were detected in different cell types throughout the follicle in pulp, bulge and the ramogenic zone. In mature barb ridges transcripts were broadly distributed throughout including barbule, marginal and axial plates, and even the feather sheath (**Figures 3C & 4, Supplementary Figure S8**). This qualitative assessment was corroborated by a quantitative measure of melanocyte-specificity S_mel_ (see Methods): a value of 0.5 indicates ubiquitous gene expression independent of cell type, larger values reflect increasing melanocyte specificity reaching 1 if expression is fully limited to melanocytes. The internal control gene ACTB showed no expression bias reflecting ubiquitous expression across all cells in the histological section of the feather follicle (median S_mel_ ACTB_S1: 0.52, ACTB_S2: 0.50). On the contrary, TYRP1, HPGDS and SLC45A2 were strongly biased towards expression in melanocytes (median S_mel_ 0.94, 0.90, 0.93 respectively). MITF showed only a slight bias towards melanocytes with a specificity index of 0.71 (**Figure 5A**). In summary, this provides evidence for melanocyte-specificity of TYRP1, HPGDS and SLC45A2, melanocyte affinity for MITF and ubiquitous expression of the ACTB control.

**Figure 4:**
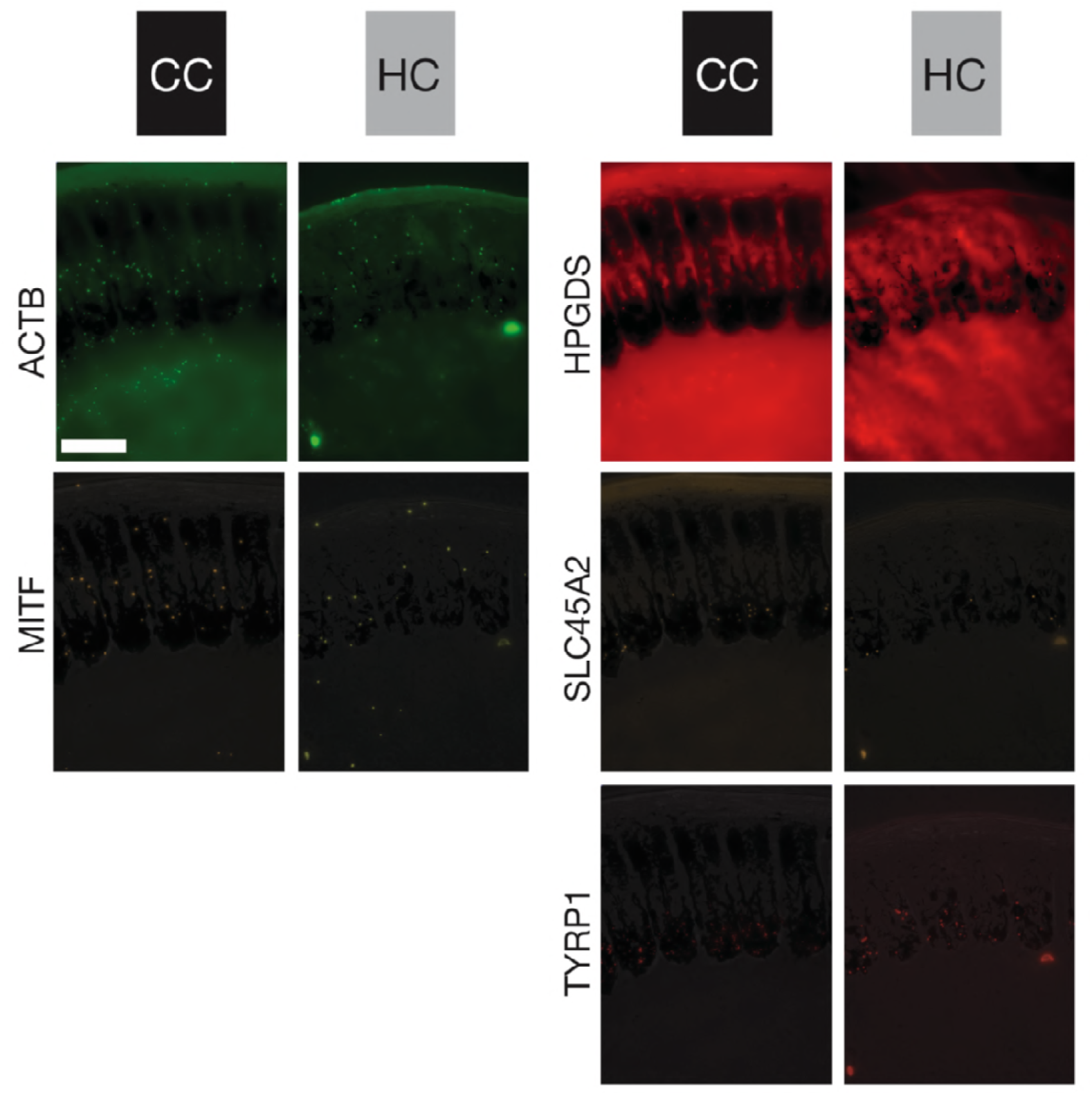
Five-way *in situ* gene expression in an explicit histological context. Illustration of *in situ* stained mRNA transcripts of five genes on the same histological cross section of carrion crow (CC black) and hooded crow (HC grey) follicles sampled from torso at 1000 μm above the dermal papilla. Bright-field images are superimposed for orientation and direct comparison among genes. The images represent one of five serial sections used for statistical analyses. ACTB and MITF (left column) are ubiquitously expressed in all cell types of a follicle. In contrast, HPGDS, SLC45A2 and TYRP1 (right column) are restricted to melanocyte cell bodies; bar = 50 μm.

**Figure 5:**
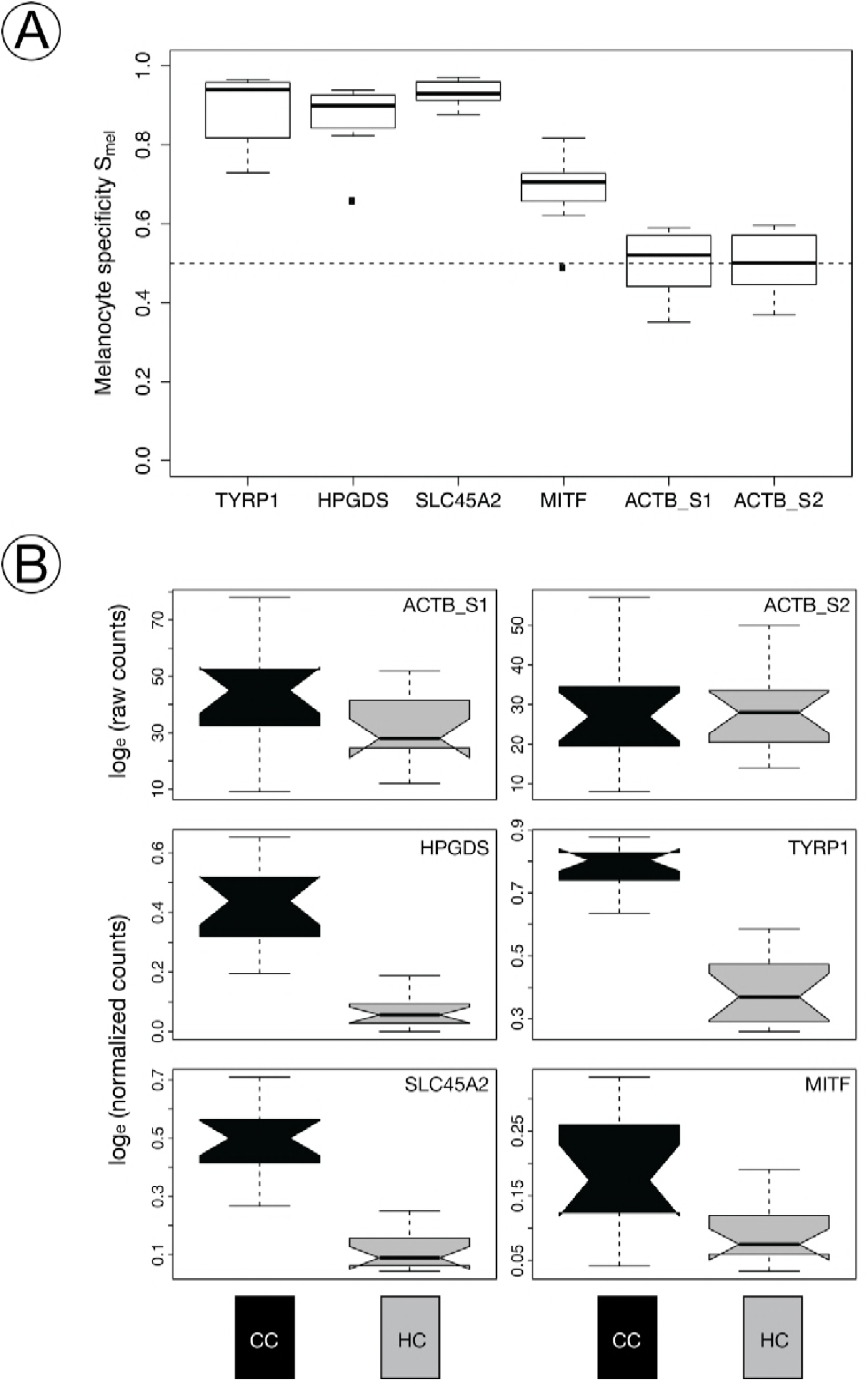
Specificity of gene expression and differentiation by taxon and pigmentation intensity. A) Boxplot of the specificity index of targeted genes. TYRP1, HPGDS, and SLC45A2 are near-exclusively expressed in melanocytes. In contrast, MITF transcripts are less specific to melanocytes, but include a variety of other cell types. **B)** Notched boxplot of raw counts (ACTB gene) or normalized *in situ* RCA signals of targeted mRNA transcripts (HPGDS, SLC45A2, TYRP1, MITF) in melanocytes at 1000 μm above the dermal papilla of carrion crow (CC) and hooded crow (HC) torso feathers. Gene expression of all four genes is significantly higher in melanocytes of carrion crow (CC black), compared to hooded crows (HC grey). For gene expression of head see **Supplementary Figure S10**.

### Gene expression transition along the proximal-distal axis of feather

We quantified the level of gene expression along the proximal-distal axis of the feather follicle (**Figure 6)**. Absolute abundance of transcripts differed, as expected, among genes and by follicle origin (black CC torso vs. grey HC torso). For the internal control gene ACTB transcript abundance was high throughout. Regardless of taxon, body region or pigmentation intensity, gene expression started at the very proximal end in the bulge region, peaked in the ramogenic zone at around 500 μm, was then stably maintained and after several millimeters slowly declined. In the highly pigmented follicles of CC torso, gene expression profiles of the melanocyte-specific TYRP1 gene differed markedly from the pattern of the ubiquitously expressed ACTB gene. mRNA abundance of this core melanosomal enzyme closely mirrored melanocyte maturation as inferred from morphology and melanin deposition (see above). The first signals of TYRP1 transcripts were detected in earliest developing melanocytes in the collared bulge. Signals quickly increased in concert with melanocyte maturation in the ramogenic zone. Expression peaked at around 1 mm from the base where spherical melanocytes were fully differentiated. Beyond this region expression gradually leveled off indicating reduced activity of melanogenesis. The other melanocyte-specific genes HPGDS and SLC45A2 had overall lower expression levels, but otherwise mirrored the mRNA abundance pattern of TYRP1 transcripts. A noteworthy difference was the near-absence at early stages of melanocyte maturation in the ramogenic zone. Expression profiles of the broadly expressed MITF gene were similar to TYRP1 with a steep onset of expression in the ramogenic zone, but with a somewhat earlier peak of expression at around 1 mm from the base of the follicle. In follicles sampled from grey torso in HC, expression patterns were generally similar to those in black torso feather despite reduced signal counts of each gene. Expression profiles in follicles sampled from head were similar in shape, but showed lower abundance for HPGDS, SLC45A2 and MITF **(Supplementary Figure S9).**

**Figure 6:**
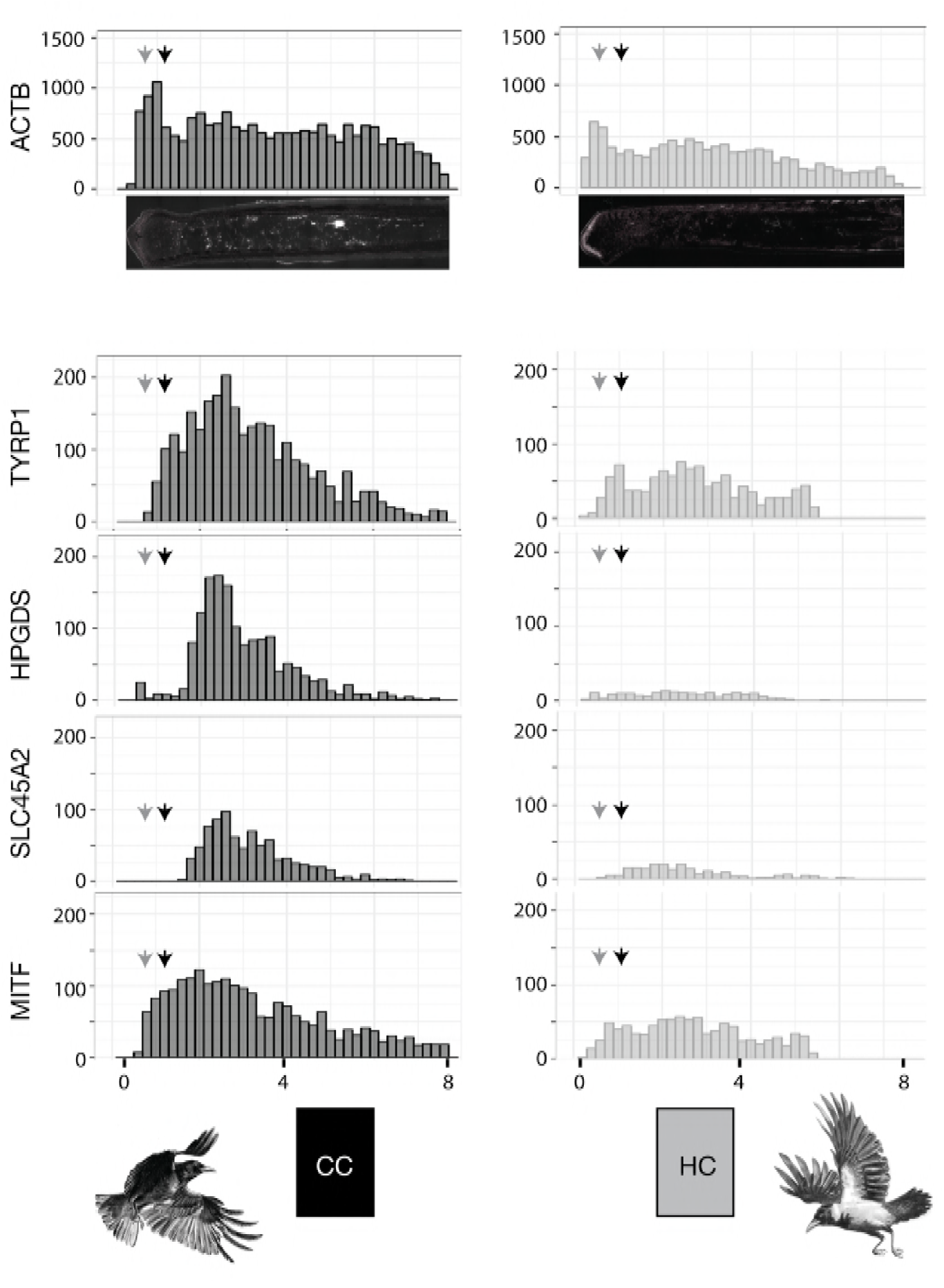
Longitudinal gene expression patterns of targeted mRNA transcripts. Histogram summarizing mRNA transcript abundance along the proximal-distal axis of feather follicles sampled from black torso of carrion crows (CC, left column) and grey torso of hooded crows (HC, right column). Each bin represents the raw counts of targeted RCA signals from mRNA transcripts of each candidate gene. Grey and black arrows indicate the position of cross sections at 500 μm and 1,000 μm above the dermal papilla, respectively. (x-axis is the distance from the proximal end in mm. y-axis is accumulated raw RCS signal counts). For gene expression pattern of follicles sampled from head see **Supplementary Figure S9**.

### Comparison of gene expression

We formally tested whether abundance of mRNA transcripts within melanocytes differed among taxa (CC vs. HC), body region (head vs. torso) and feather pigmentation (black vs. grey). mRNA transcripts were quantified in the cell body of melanocytes for all four target genes and ACTB as internal control. Generalized linear mixed effect models using statistical model selection based on the Akaike Information Criterion suggested an effect of taxon and pigmentation for TYRP1, HPGDS and MITF, and an additional effect of body region for SLC45A2 (**Table 1**). Overall, the parameter with the by far strongest influence was feather pigmentation (wAIC = 1.00 for all genes) followed by taxon (wAIC ranging from 0.57 for TYRP1 to 0.83 for MITF). Gene expression was slightly higher in head of hooded crows (black in both species), but substantially lower in follicles sampled from the grey torso region (**Figure 5B, Supplementary Figure S10**). These results were independent of melanocyte maturity both at 500 μm vs. 1000 μm above the dermal papilla.

**Table 1:**
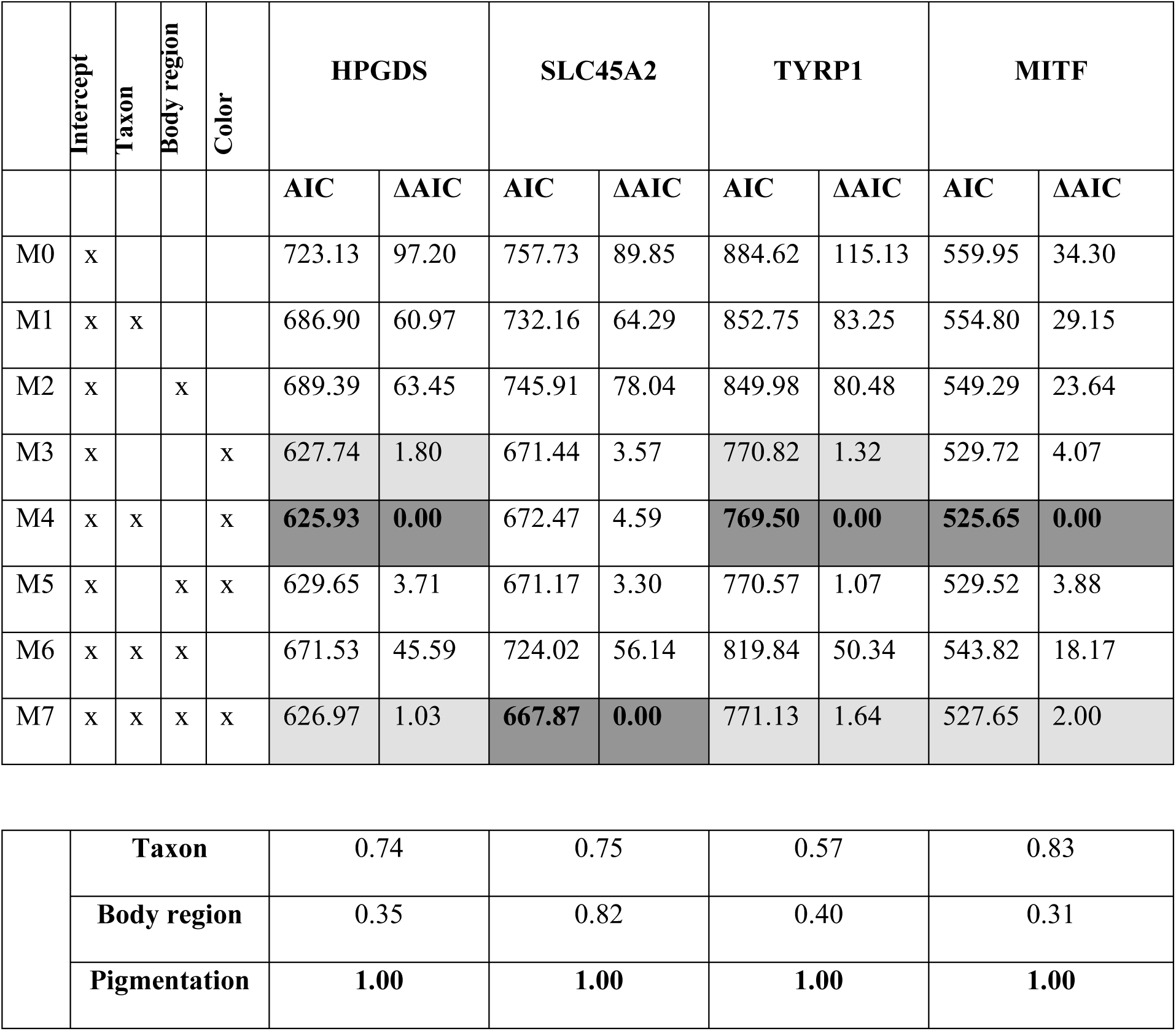
Summary of the statistical models exploring the effect of taxon (carrion vs. hooded crow), body region (head vs. torso) and color (black vs. grey) on normalized gene expression of the four target genes. *upper part*: Factors included in each model are indicated by ‘x’ (e.g. M7: gene expression ∼ intercept + taxon + bodyregion + color). Akaike’s information criterion (AIC) and difference to the best model (δAIC) are shown. The best model is shown in bold on dark grey background; other models with support (δAIC≤2) are highlighted in light grey. *lower part*: Akaike’s weights (wAIC) indicating the relative importance of a factor are shown for each gene.

|  | Intercept | Taxon | Body region | Color | HPGDS |  | SLC45A2 |  | TYRP1 |  | MITF |  |
| --- | --- | --- | --- | --- | --- | --- | --- | --- | --- | --- | --- | --- |
|  |  |  |  |  | AIC | ΔAIC | AIC | ΔAIC | AIC | ΔAIC | AIC | ΔAIC |
| M0 | x |  |  |  | 723.13 | 97.20 | 757.73 | 89.85 | 884.62 | 115.13 | 559.95 | 34.30 |
| M1 | x | x |  |  | 686.90 | 60.97 | 732.16 | 64.29 | 852.75 | 83.25 | 554.80 | 29.15 |
| M2 | x |  | x |  | 689.39 | 63.45 | 745.91 | 78.04 | 849.98 | 80.48 | 549.29 | 23.64 |
| M3 | x |  |  | x | 627.74 | 1.80 | 671.44 | 3.57 | 770.82 | 1.32 | 529.72 | 4.07 |
| M4 | x | x |  | x | **625.93** | **0.00** | 672.47 | 4.59 | **769.50** | **0.00** | **525.65** | **0.00** |
| M5 | x |  | x | x | 629.65 | 3.71 | 671.17 | 3.30 | 770.57 | 1.07 | 529.52 | 3.88 |
| M6 | x | x | x |  | 671.53 | 45.59 | 724.02 | 56.14 | 819.84 | 50.34 | 543.82 | 18.17 |
| M7 | x | x | x | x | 626.97 | 1.03 | **667.87** | **0.00** | 771.13 | 1.64 | 527.65 | 2.00 |

TABLE 1
|  |  |  |  |  |  |
| --- | --- | --- | --- | --- | --- |
|  | **Taxon** | 0.74 | 0.75 | 0.57 | 0.83 |
|  | **Body region** | 0.35 | 0.82 | 0.40 | 0.31 |
|  | **Pigmentation** | **1.00** | **1.00** | **1.00** | **1.00** |

## DISCUSSION

### Functional validation of genes by padlock probe combined with rolling circle amplification

The study of animal coloration has importantly contributed to our understanding of the genetic basis underlying inheritance and evolution (8, 45). Long confined to traditional model organisms such as mouse or fruit fly high-throughput nano-sequencing technology now allows investigating the genetic basis of color variation across a broad array of organisms (21). Yet, functional genetic studies remain difficult for the vast majority of species that are not amenable to assays such as gene knock-outs, RNA interference or CRISPR-Cas9 modification that are standard for laboratory models (46).

We here suggest a versatile approach for functional validation of genetic signals from genome scans or association mapping applicable to a wide variety of natural systems. In contrast to bulk mRNA sequencing which integrates over diverse cell populations, *in situ* padlock probes are a flexible tool to visualize mRNA transcripts of any gene of interest in a the native, histological context (47). Combined with rolling circle amplification single mRNA molecules are imaged as bright, round-shaped fluorescent signals that can readily be quantified using automated image analyses. In contrast to conventional *in situ* hybridization relying on relative fluorescence intensity, the approach chosen here results in numeric count data. This property allows for flexible statistical analyses of complex treatment design. Moreover, referencing counts to a control gene known to be unaffected by the contrast in question (housekeeping gene) controls for methodological variation in staining efficiency and allows integrating data across multiple sections into one statistical analysis.

### Single mRNA transcript counts are overall well correlated with RNA-seq based quantification

Using an evolutionary model for incipient speciation characterized by a striking polymorphism in feather pigmentation, we demonstrate the utility of the approach to characterize gene expression relevant to natural phenotypic variation. Padlock probes for detection of mRNA was recently developed (47) and has so far only been used in a few studies in a biomedical context (48–50). Following the guidelines published (29) we designed probes for five different targets and applied them to analyze expression profiles in the complex tissue of nascent feather follicles. Given the added difficulty of working with keratizined, water repelling feather material suggests that the method is applicable across a wide variety of tissues. Relative estimates of steady-state mRNA concentration as measured by FPKM (fragments per kilobase per million reads) via bulk mRNAseq in skin tissue containing re-growing feather follicles differed by more than tenfold among candidate genes. SLC45A2 and MITF were most lowly expressed, followed by HPGDS with intermediate levels of expression and highest values for TYPR1 (see Fig. 4 in Poelstra et al., 26). The normalized mRNA counts derived here using the padlock/RCA assay were highly correlated with the FPKM based estimates in all cells (R^2^=0.92, p<0.001), and to a lower extent when exclusively considering melanocyte-specific transcripts (R^2^=0.79, p<0.001). The correlation was largely unaffected by body region and taxon. This suggests that the method is not only suited to compare expression by treatment separately for each gene, but also to relate absolute transcript numbers from different genes normalized by the same control.

### Cell-type specific quantification of the ubiquitously expressed MITF transcripts reveals taxon-specific expression

Poelstra et al. (26) found widespread differential expression in the melanogenesis pathway including genes such as SLC45A2 or TYRP1 which are well characterized to act downstream in melanin production (51). MITF, however, which assumes a central regulatory function in melanogensis by integrating signals from several interacting pathways (WNT, cAMP-dependent, MAPK) was not differentially expressed. Using the padlock/RCA approach this conflict could be resolved. Developing a measure of cell-type specificity we could show that transcripts of SLC45A2 and TYRP1 were near-exclusively restricted to melanocyte where they also were enriched in the cell body relative to dendrites. MITF transcripts, on the contrary, were ubiquitously expressed. Considering melanocyte-specific transcripts alone MITF expression followed the expected pattern of significant upregulation in black carrion crow feathers (26). The lack of differential expression in MITF using bulk mRNA sequencing was thus not due to a different functional role in avian regulatory networks, but reflects signal dilution by other cell types. A central regulatory role of MITF in melanogenesis is therefore likely to apply to avian systems in general, and specifically to divergence pigmentation patterns between hooded and carrion crows.

MITF has been shown to modulate cell-cycle progression altering melanocyte survival and turnover and to regulate the amount of melanin produced in melanocytes (51, 52). Our data lend support the latter function. Pigmentation differences were exclusively due to change in the concentration of eumelanin contained in rod-shaped granules in both taxa, and there was no evidence for differences in melanocyte shape or development. Moreover, while absolute transcript numbers differed significantly between grey hooded crow feather follicles and black carrion crow follicles, longitudinal expression patterns did not. In both taxa MITF transcripts were detected at the very onset of melanocyte development and preceded expression of TYPR1 and SLC45A2. This is consistent with a role of MITF in early melanocyte development and in regulation of downstream elements such as TYPR1 and SLC45A2 being transcribed at a more advanced stage of melanocyte maturation. Altogether, this suggests a role of MITF in regulating differential melanin concentration among taxa.

Using population genomic screens in combination with bulk mRNA sequencing it had been suggested in the crow system that MITF may itself be regulated by upstream elements that have diverged during the course of evolution (26, 28). These include CACNG calcium channel genes regulating AMPA receptors known to influence expression of MITF.

### A functional role for the HPGDS gene in melanogenesis

Another divergent candidate was HPGDS, a gene that is generally associated with gluthathione metabolism with no clear bearing on melanogensis. Yet, HPGDS may interact with MITF through SOX9 or by its calcium ion binding functionality (26 and references therein, 28). The observation of this study that HPGDS transcripts were limited to melanocytes lends further support to a potential role in melanogensis. Moreover, congruent with MITF, it was expressed at an early stage of melanocyte maturation which is consistent with the proposition that it may be involved in regulatory feed-back with MITF.

## AUTHOR CONTRIBUTIONS

CW and JW conceived of the study and together with OS designed the experiments with input from AK in primer design and PR in image analysis. During a pilot phase RM developed parts of the methodological set up. JW collected the samples and together with JB was responsible for animal husbandry. CW and JW wrote the manuscript with input from all other authors.

## ACKNOWLEDGEMENTS

We express our gratitude to Sven Jakobsson for providing the infrastructure for animal husbandry at Tovetorp research station. We would also like to thank Christen Bossu, Jelmer Poelstra and Matthias Weissensteiner for their contribution in obtaining samples. Kristaps Solokovskis, Thomas Giegold, Nils Andbjer, TamaraVolkmer, Barbara Martinschitsch and Luisa Sontheimer were of invaluable support for raising and maintaining the captive crow population. Martin Wikelski, Inge Müller and additional staff from the Max-Planck-Institute for Ornithology in Radolfzell facilitated sampling in

Germany and transport to Sweden. Margareta Mattson and Helena Malmikumpu provided help with sectioning.

